# An intracellular tripeptide Arg-His-Trp of serum origin detected in MCF-7 cells is a possible agonist to β2 adrenoceptor

**DOI:** 10.1101/2020.12.28.424618

**Authors:** Hritik Chandore, Ajay Kumar Raj, Kiran Bharat Lokhande, K. Venkateswara Swamy, Jayanta K. Pal, Nilesh Kumar Sharma

**Author notes:** MIT-School of Bioengineering Sciences & Research, MIT-Art, Design and Technology University, Pune, Maharashtra, India, 412201. Corresponding author: Dr. Nilesh Kumar Sharma, Professor, Cancer and Translational Research Lab, Department of Biotechnology, Dr. D. Y. Patil Biotechnology & Bioinformatics Institute, Pune, Dr. D. Y Patil Vidyapeeth Pune, Pune, MH, 411033.

## Abstract

**Background:** The need of agonists and antagonists of β2 adrenoceptor (β2AR) is warranted in various human disease conditions including cancer, cardiovascular and other metabolic disorders. However, the sources of agonists of β2AR are diverse in nature. Interestingly, there is a complete gap in the exploration of agonists of β2AR from serum that is a well-known component of culture media which supports growth and proliferation of normal and cancer cells in vitro.

**Methods:** In this paper, we employed a novel vertical tube gel electrophoresis (VTGE)-assisted purification of intracellular metabolites of MCF-7 cells grown *in vitro* in complete media with fetal bovine serum (FBS). Intracellular metabolites of MCF-7 cells were then analyzed by LC-HRMS. Identified intracellular tripeptides of FBS origin were evaluated for their molecular interactions with various extracellular and intracellular receptors including β2AR (PDB ID: 2RH1) by employing molecular docking and molecular dynamics simulations (MDS). A known agonist of β2AR, isoproterenol was used as a positive control in molecular docking and MDS analyses

**Results:** We report here identification of a few novel intracellular tripeptides, namely Arg-His-Trp, (PubChem CID-145453842), Pro-Ile-Glu, (PubChem CID-145457492), Cys-Gln-Gln, (PubChem CID-71471965), Glu-Glu-Lys, (PubChem CID-11441068) and Gly-Cys-Leu (PubChem CID145455600) of FBS origin in MCF-7 cells. Molecular docking and MDS analyses revealed that among these molecules, the tripeptide Arg-His-Trp shows a favorable binding affinity with β2AR (−9.8 Kcal/mol). Furthermore, agonistic effect of this tripeptide, Arg-His-Trp is significant and comparable with that of a known agonist of β2AR, isoproterenol.

**Conclusion:** In conclusion, we identified a unique Arg-His-Trp tripeptide of FBS origin in MCF-7 cells by employing a novel approach. This unique tripeptide Arg-His-Trp is suggested to be a potential agonist of β2AR and it may have applications in the context of various human diseases like bronchial asthma and chronic obstructive pulmonary disease (COPD).

**NOVELTY & IMPACT STATEMENTS:**

- This paper reports on a novel vertical tube gel electrophoresis (VTGE) system that assisted in the purification and identification of a few intracellular tripeptides in the in vitro grown breast cancer cells, MCF-7ls.
- Molecular docking and molecular dynamics simulation analyses strongly suggest that the tripeptide Arg-His-Trp among others forms the most stable ligand-protein complex with β2 adrenoceptor (β2AR). Its binding affinity and the nature of molecular interactions are comparable or even better than the known agonists of β2AR.
- This tripeptide Arg-His-Trp is predicted to show manyfold less cytotoxicity, mutagenicity, cardiotoxicity, drug-drug interactions, microsomal stability, and drug-induced liver injury over the other known agonists of β2AR.
- The tripeptide Arg-His-Trp is therefore suggested as an effective agonist of β2AR and this may be validated in future, in preclinical and clinical models.

## Introduction

Human cells are endowed with several thousand genes that encode various proteins including receptors that contribute to cellular growth and proliferation of normal and cancer cells (Vander Heiden et al., 2009; Tennant et al., 20101). These growth receptors require ligands like growth factors, peptides and other chemical factors from their surroundings and growing environments including culture medium in case of in vitro cell culture (Sullivan et al., 2016; Li et al., 2019). However due to lack of methods and processes to study intracellular metabolites including tripeptides, the role of such molecules naturally present in the culture medium enriched with fetal bovine serum (FBS, is not known.

It is well-known that FBS is used as the key source of various molecules for supporting growth of normal and cancer cells (Wu et al., 2008; Ulanovskaya et al., 2013; Kraus et al., 2014; Crujeiras et al., 2018). Furthermore, tripeptides as biological agents from natural sources including FBS or synthetic sources are reported to have an extent of various activities/properties such as antioxidative, pro-growth, differentiation, tissue repair and various other activities under normal physiological conditions (Sensenbrenner et al., 1975; Pickart and Thaler, 1979; Congote, 1986; Hsu et al., 2005; Turpeinen et al., 2011; Xiao et al., 2012; Fang et al., 2017; Daskalaki et al., 2018). However, an efficient method and process to purify the biologically active small peptides and their compositions from the FBS is not available. Thus, there appears to be a limitation in our knowledge on the chemical nature of FBS in the form of tripeptides that enter the growing cells, their identifications and also on their potential biological roles.

The biological effects of these small molecules are known to be mediated through activation of various molecular cell signaling pathways and molecules including β2 adrenoceptor (β2AR), a type of G-protein coupled receptor that is known to support the growth and activation of cells (Vandewalle et al., 1990; Pérez Piñero et al., 2012; Szpunar et al., 2013; Liu et al., 2015; Kim et al., 2016; Castillo et al., 2017; Gargiulo et al., 2017; He et al., 2017; Clément-Demange et al., Gargiulo et al., 2020; Zhou et al., 2020). At the same time, β2AR is reported to be defective and dysfunctional in several human disorders including asthma, chronic obstructive pulmonary disease (COPD), neurodegenerative diseases, and immune disorders (Ahuja and Smith, 2007; Audet and Bouvier, 2008; Katritch et al., 2009; Dror et al., 2011; Kahsai et al., 2011; Rasmussen et al., 2011; Rasmussen et al., 2011; Rosenbaum et al., 2011; Vanni et al., 2011; Lebon et al., 2012).

In the literature, agonists and antagonists have been reported that modulate the activity of β2AR in various human diseases including metabolic disorders, cancers, and cardiovascular diseases (Wieland et al., 1996; Cherezov et al., 2007; Audet and Bouvier, 2008; Katritch et al., 2009; Dror et al., 2011; Kahsai et al., 2011; Rasmussen et al., 2011; Rasmussen et al., 2011; Rosenbaum et al., 2011; Vanni et al., 2011; Lebon et al., 2012; Weiss et al., 2013; Dilcan et al., 2013; Weichert et al., 2014; Zhang et al., 2020). Interestingly, the effects of agonists and activators of β2AR are known to induce growth, proliferation and invasion of ER positive breast cancer cells including MCF-7 cells (Vandewalle et al., 1990; Pérez Piñero et al., 2012; Szpunar et al., 2013; Liu et al., 2015; Kim et al., 2016; Castillo et al., 2017; Gargiulo et al., 2017; He et al., 2017; Clément-Demange et al., Gargiulo et al., 2020; Zhou et al., 2020). However, tripeptides, particularly of FBS origin, as agonists of β2AR that supports the growth and activation of cells is not known. Hence, our rationale to explore tripeptides from FBS that support the growth of in vitro grown cells and to explore their agonistic pro-growth effects on the potential receptor β2AR, is valid.

We report here on a novel approach to detect biologically important tripeptides from FBS that is commonly used for *in vitro* growth and maintenance of normal, cancer and other types of cells. Purification of such intracellular tripeptides has been assisted by a novel methodology using vertical tube gel electrophoresis (VTGE) system. Furthermore, evaluation of their agonistic effects against β2AR has been carried out by molecular docking and MDS during 10ns to understand the stability of the peptides within the binding cavity of β2AR receptor.

## Materials and methods

### Trypan blue dye exclusion assay

The MCF-7 cancer cells were grown to 60-70% confluency and cells were harvested and plated onto six well plates at the seeding density of 150,000 cells per well. In the next day, the cell-plated wells were organized in triplicates and were added with complete Dulbecco’s Modified Eagle Medium (DMEM, HiMedia Laboratories Pvt. Ltd. Mumbai, Catalogue No: ALA007A) containing 10% FBS (Gibco Model Catalogue No 26140087) and 1% Penicillin and streptomycin antibiotics with 0.5% DMSO. At the end of 48 hr of growth at 37°C in CO2 incubator, cells were harvested and collected by routine method of Trypsin-EDTA assisted trypsinization and centrifugation. A routine Trypan dye exclusion assay was performed to determine the viable cells.

In this assay, MCF-7 cells were grown in an optimal condition including presence of FBS and added with 0.5% DMSO that is mostly used as a solvent control in most of the drug treatment assays.

### Dual ethidium bromide/acridine orange fluorescent staining

For this assay, MCF-7 cancer cells were grown for 48 hr in complete Dulbecco’s Modified Eagle Medium (DMEM, HiMedia Laboratories Pvt. Ltd. Mumbai, Catalogue No: ALA007A) containing 10% FBS (Gibco Model Catalogue No 26140087) and 1% Penicillin and Streptomycin antibiotics with 0.5% DMSO. After 48 hr, routine trypsinization was done to harvest cells and suspension of cells were prepared. Next, prepared MCF-7 cancer cell suspension (10 μl) was transferred to glass slides. From the stock of dual fluorescent staining solution (100 μg/ml AO and 100 μg/ml EB), one μl was dropped onto glass slides with cell suspension and covered with a coverslip. Then immediately, MCCF-7 cancer cells were examined for viability and cell death using a fluorescent microscope (OLYMPUS, Japan) (Kumar et al., 2020).

### Preparation of intracellular lysate

At the end of 48 hr of growth of MCF-7 cancer cells, standard protocol and washing procedure were done appropriately to exclude the possibility of external media contamination. For the preparation of intracellular lysate of one million cells, 500 μl of hypotonic extraction buffer (NaCl 20 mM and KCl 2.7 mM) was used to lyse MCF-7 cancer cells by employing a 31-gauge needle and dounce glass homogenizer to achieve mechanical disruption of cells. Next, lysed MCF-7 cancer cells were centrifuged at 12000 g for a duration of 30 min to obtain clear cell lysate of MCF-7 cancer cells. In this paper, “Whole Cell Lysate” of MCF-7 cancer cells was hypotonically prepared and used for the preparation of novel intracellular tripeptides by employing a novel VTGE purification tool (Kumar et al., 2020).

### Purification of intracellular metabolites by VTGE

Finally, 250 μl of whole cell lysate was mixed with 500 μl of hypotonic extraction buffer. Thereafter, the prepared whole cell lysate was mixed with 250 μl of 4X loading buffer (0.5 M Tris, pH 6.8 and Glycerol. The above prepared whole cell lysate was loaded on the VTGE purification system (Figure S1) with a matrix of 15% acrylamide gel (acrylamide: bisacrylamide, 30:1). The fractionated metabolites were eluted in the 5X running buffer (960 mM glycine of pH at 8.3). During purification of intracellular metabolites, 1500-2500 mW of power was maintained for 90-120 minute. Finally, these collected intracellular metabolites as elutes in the elution buffer were stored at −20°C for further LC-HRMS based identification and analysis (Sharma et al., 2019, Kumar et al., 2020). Here, it is important to mention that eluted intracellular metabolites in the used elution buffer is quite stable at room temperature during transportation and handling for LC-HRMS based analysis.

### Identification of intracellular metabolites by LC-HRMS

The eluted intracellular metabolites containing novel tripeptides of FBS were identified by LC-HRMS analytical system. The analysis of metabolites employed Agilent TOF/Q-TOF Mass Spectrometer station Dual AJS ESI ion source. During this process, RPC18 Hypersil GOLD C18 100 x 2.1 mm-3 μm was used as LC column. Mass spectrometry was performed in a positive mode. During LC separation, mobile phase of 100% Water (0.1% FA in water) and 100% Acetonitrile (90% ACN +10% H_2_O+ 0.1% FA) was used in the form of 95% and 5% proportion. Next, these identified novel intracellular tripeptides namely Arg-His-Trp, Pro-Ile-Glu, Cys-Gln-Gln, Glu-Glu-Lys and Gly-Cys-Leu were submitted to EMBL-ChEBI and NCBI PubChem so that curation of structure and generation of sdf file format was performed (Sharma et al., 2019).

### Molecular Docking

These novel intracellular tripeptides Arg-His-Trp, (PubChem CID-145453842), Pro-Ile-Glu, (PubChem CID-145457492), Cys-Gln-Gln, (PubChem CID-71471965), Glu-Glu-Lys, (PubChem CID-11441068) and Gly-Cys-Leu (PubChem CID-145455600) were downloaded from ChEBI database. The molecular editor software Avogadro (Morris et al., 2009) was used to energy minimization of retrieved peptides to get energetically stable conformation and minimized conformations were converted into PDB file format for the further calculations. Protein Data Bank (PDB) was used to download the crystal structure of β2AR (PDB ID: 2RH1) (https://www.rcsb.org). Then, β2AR was subjected to the AutoDock Tool v4.2.1 to prepare the protein for docking procedure along with ligand as tripeptide. AutoDock Vina Software was employed to study molecular docking of novel intracellular tripeptides Arg-His-Trp, Pro-Ile-Glu, Cys-Gln-Gln, Glu-Glu-Lys, Gly-Cys-Leu and a known agonist Isoproterenol (PubChem CID:5808) with β2AR (Trott and Olson, 2010). After the successful molecular docking, binding conformations pattern of these novel tripeptides were analyzed through discovery studio visualizer (DSV3, 2020).

### Molecular Dynamics Simulations

The molecular dynamics simulation was performed up to 10ns for a novel tripeptide Arg-His-Trp with highest binding affinity towards β2AR receptor using Desmond software (Schrödinger, 2019) to assess the binding strength of the control and screened peptides with cell surface adrenergic receptor. Using a system builder of Desmond in the Maestro program, the complex system was immersed in a water-filled (TIP3P water model) orthorhombic box of 10 Å spacing. The total charge of the solvent system was neutralized by adding 2 sodium (Na+) of concentration 4.066 mM. These studies were carried out with a run of 10ns with a constant temperature of 300K using NPT (N=number of atoms, P=Pressure, T=Temperature) ensemble and OPLS-2005 force field, considering certain parameters including Martyna-Tobias-Klein barostat method, Nose Hoover chain thermostat method, integrator as MD, and constraints set as all-bonds. The conformational changes of β2AR receptor upon binding of Arg-His-Trp was recorded by using the 1000 trajectory frames generated during the 10ns MD simulation.

### Statistical analysis

Data are presented as the mean ± SD of at least three independent experiments. Differences are considered statistically significant at P < 0.05, using a Student’s t-test.

## Results

### Identifications of FBS derived intracellular tripeptides in MCF-7 cells

In view of the existing evidence on the growth promoting factors in FBS, a novel approach was employed to look into the FBS components that are implicated in cellular responses. In this direction, MCF-7 breast cancer cells were taken as model in vitro cells grown in the presence of complete culture medium supported with FBS.

Based on the simple and routinely used Trypan blue dye exclusion assay, it was ascertained that MCF-7 breast cancer cells were in optimal growth and proliferation mode in the presence of complete medium with FBS. It is important to mention that during a normal and controlled growth conditions, routinely used DMSO solvent was used to make the relevance that most of in vitro experimental work, normal growth conditions also contained solvent control including DMSO. As expected, MCF-7 breast cancer cells showed optimal growth and viability in the presence of complete media and FBS components (Figure 1 A and B). Furthermore, acridine orange and ethidium bromide dual staining assay showed expected viability up to 90% and a very low presence of non-viable cells that may be due to DMSO effects and loss of viability during the experimentation conditions (Figure 2A and B). In fact, observed growth and viability of MCF-7 cancer cells are not surprising and lack of novelty. However, an intriguing question prompted us to known if any tripeptide and similar small molecular weight compounds from FBS that can promote the growth and viability of MCF-7 cancer cells. A major hurdle was to study intracellular metabolites of growing MCF-7 cancer cells. A recent novel methodology from our lab is reported as VTGE that assists in the purification of small molecular weight compounds less than 1000 Da. We know that tripeptides are also in the range of M.W. 300 to 600 Da. Therefore, we attempted to explore the nature of FBS components those are small molecular weight compounds including tripeptides by employing a novel VTGE (Figure S2) assisted purification of intracellular components of MCF-7 cancer cells growing in the presence FBS.

**Figure 1.**
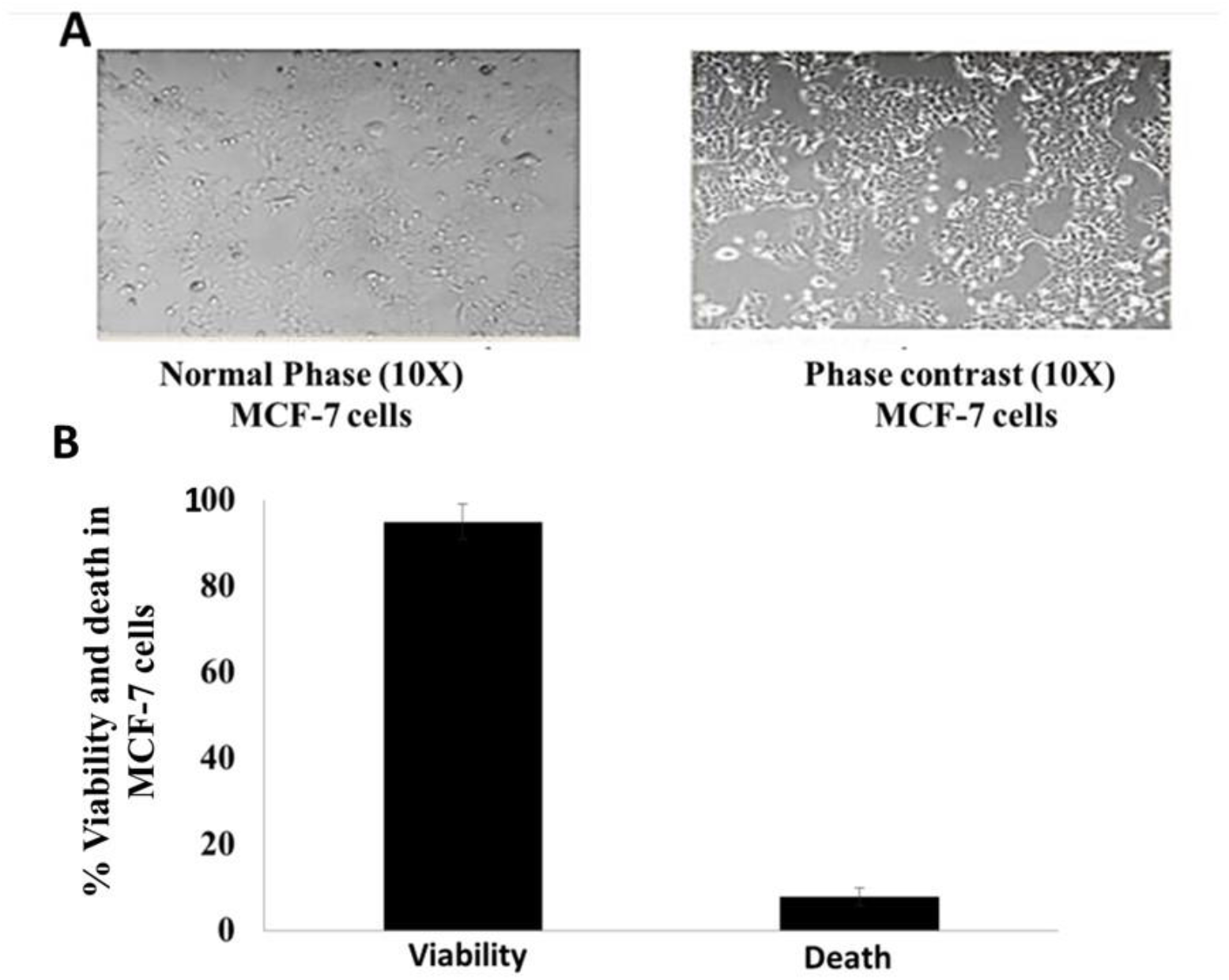
MCF-7 cancer cells show optimal growth and proliferation in the presence of complete DMEM media with FBS. Cells were grown for 48 hr followed by microscopy and Trypan blue dye exclusion assay to estimate viable and dead cells. (A) Bright light and phase contrast microscopy photographs (100 X magnification) of MCF-7 cancer cells. (B) Bar graph showing percentage viability and death of cells as determined by Trypan blue dye exclusion assay.

**Figure 2.**
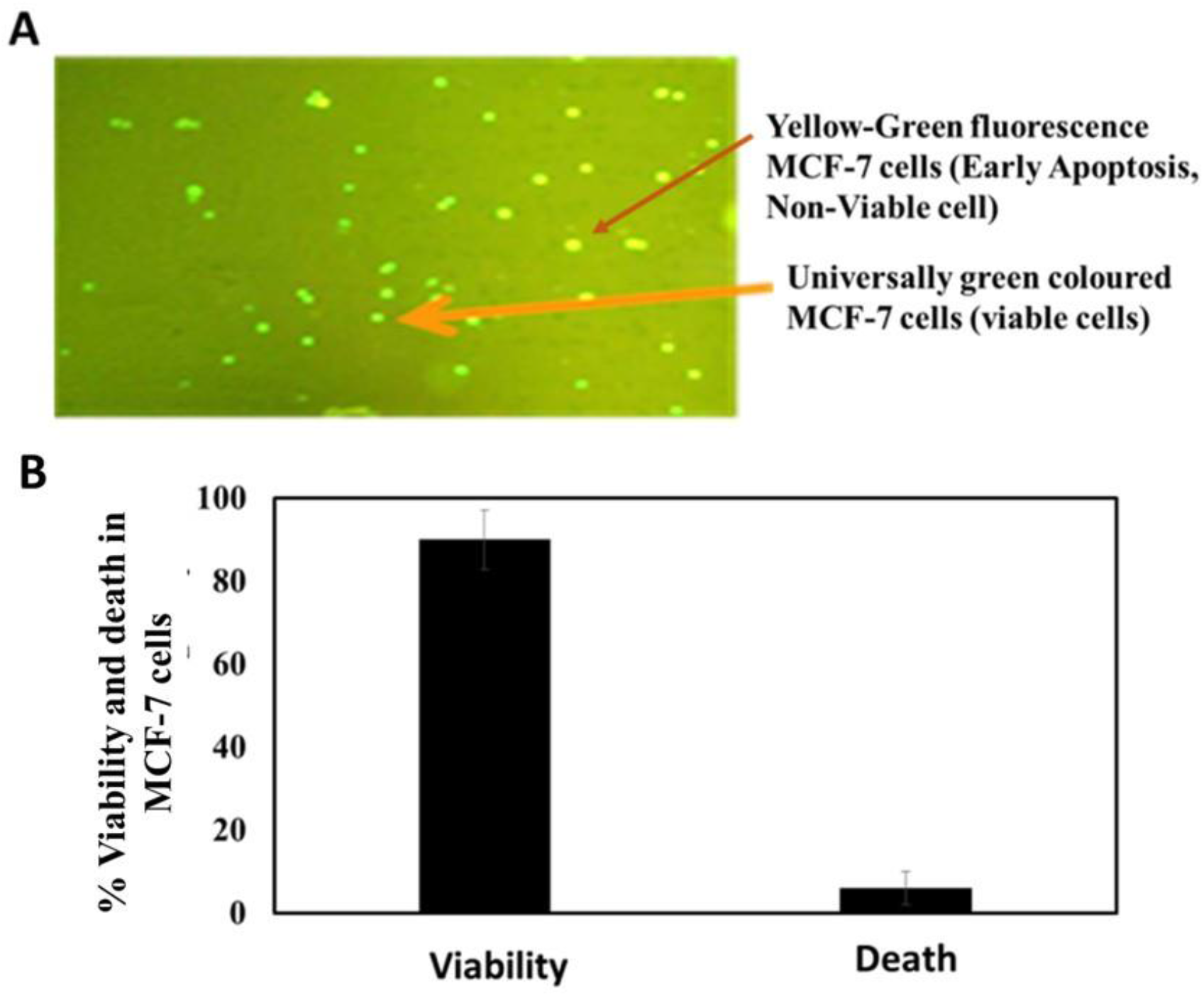
MCF-7 cancer cells show viable proliferation in the presence of complete DMEM medium with FBS. Cells were stained by using ethidium bromide and acridine orange followed by dual staining fluorescent microscopy assay. (A) Fluorescence microscopy photographs were taken at 100X magnification for MCF-7 cancer cells. Here, universally green colored MCF-7 cancer cells stained by acridine orange as viable cells and yellow-green stained MCF-7 cancer cells as stained by both acridine orange and ethidium bromide denoting early apoptotic cells. (B) Bar graph denotes the percent viability and death in MCF-7 cancer cells.

Here, novel intracellular tripeptides derived of FBS are detected in MCF-7 cancer cells and their positive ESI MS ion spectrum is presented. These tripeptides were Arg-His-Trp (Figure 3), Pro-Ile-Glu (Figure S2), Cys-Gln-Gln (Figure S3), Glu-Glu-Lys (Figure S4) and Gly-Cys-Leu (Figure S5). These intracellular tripeptides namely Arg His Trp (m/z 480.2387, mass 497.2429 and chemical formula C23 H31 N9 O4), Pro Ile Glu (m/z 340.1861, mass 357.1895 and chemical formula C16 H27 N3 O6), Cys-Gln-Gln (m/z 400.1229, mass 377.1340 and C13 H23 N5 O6 S), Glu-Glu-Lys (m/z 387.1874, mass 404.1908 and chemical formula C16 H28 N4 O8) and Gly-Cys-Leu (m/z 274.1240, mass 291.1274 and chemical formula C11 H21 N3 O4 S) showed characteristic MS ion properties. These identified tripeptides were curated for chemical structure and their ChEBI-IDs and PubChem CIDs were published (Table 1). Furthermore, these intracellular tripeptides were used for molecular docking and MDS analyses against various extracellular and intracellular target proteins.

**Figure 3.**
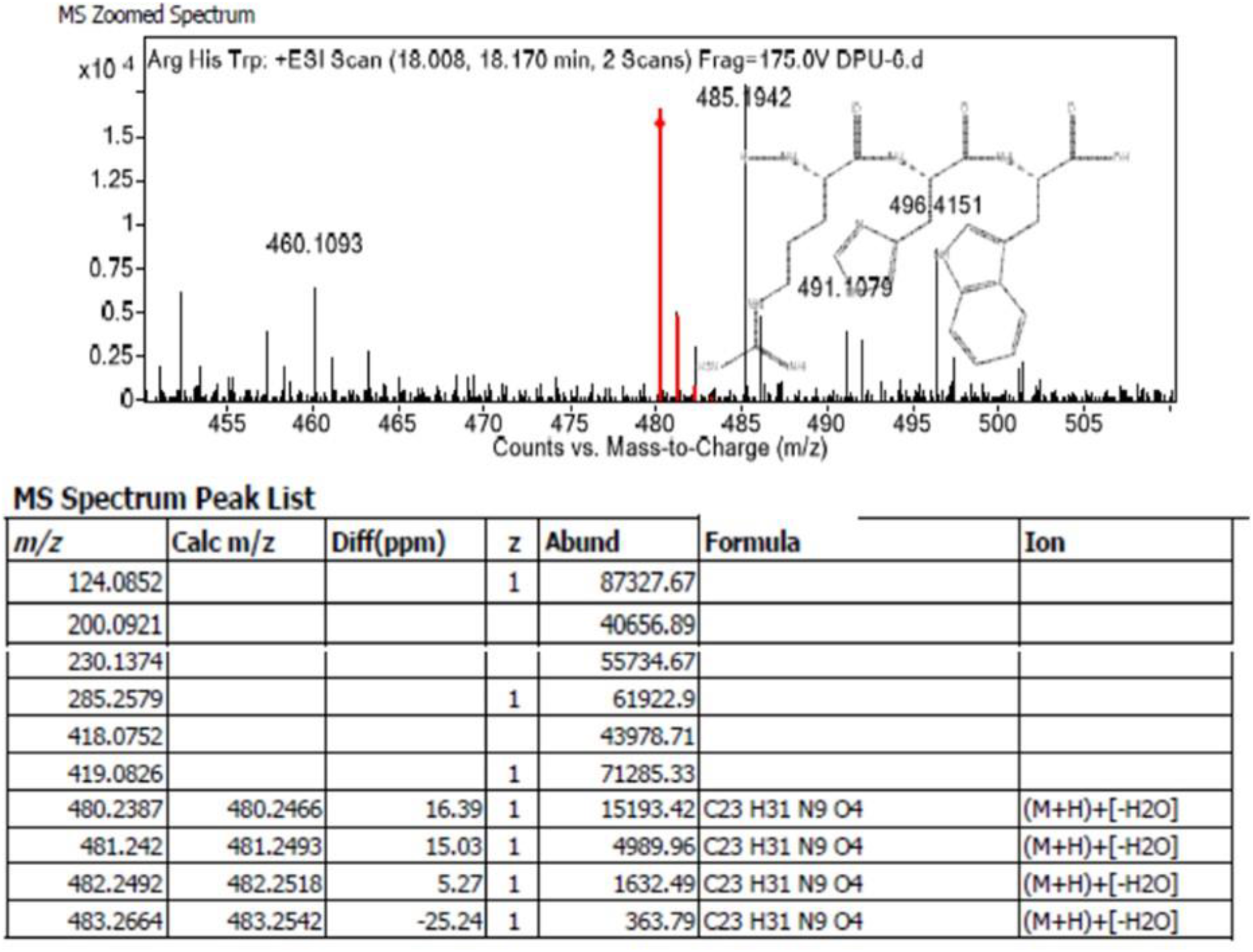
Intracellular tripeptides of FBS origin is detected in MCF-7 cancer cells. A positive ESI MS spectra and observed peak lists of intracellular triepetides Arg-His-Trp detected in MCF-7 cancer cells.

**Table 1:**
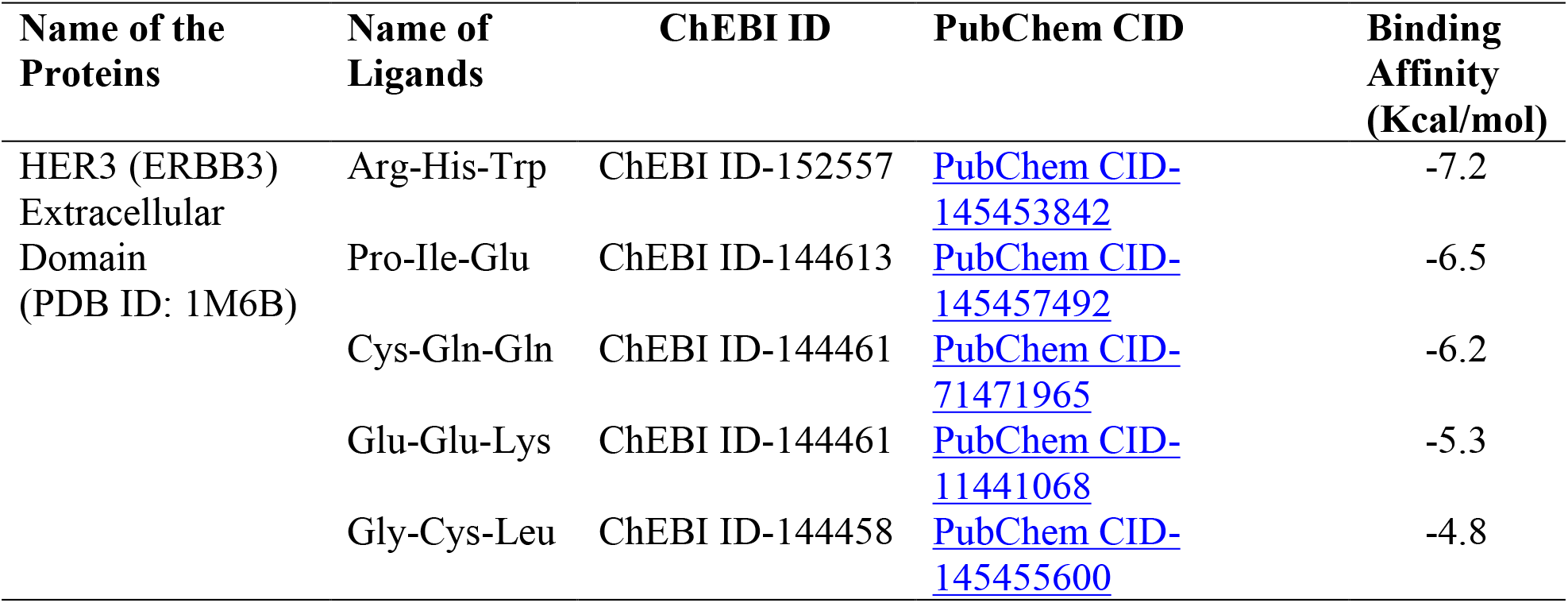
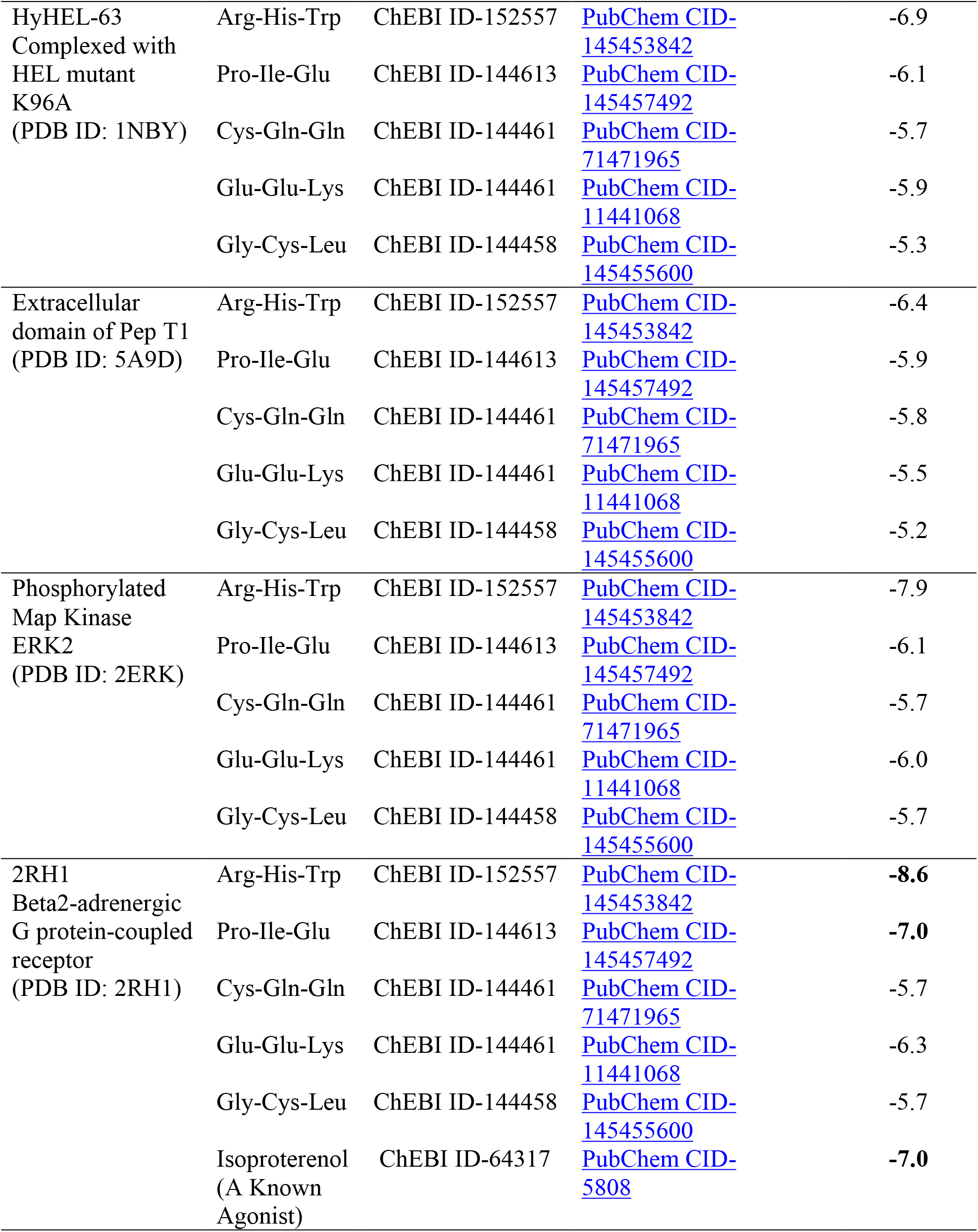
The details on screening of binding affinities of identified tripeptides from FBS with extracellular pro-growth receptors and selected intracellular proteins. Here, molecular interactions between tripeptides and target proteins are evaluated by using AutoDock Vina blind docking tool.

| Name of the Proteins | Name of Ligands | ChEBI ID | PubChem CID | Binding Affinity (Kcal/mol) |
| --- | --- | --- | --- | --- |
| HER3 (ERBB3) Extracellular Domain (PDB ID: 1M6B) | Arg-His-Trp | ChEBI ID-152557 | PubChem CID-145453842 | -7.2 |
|  | Pro-Ile-Glu | ChEBI ID-144613 | PubChem CID-145457492 | -6.5 |
|  | Cys-Gln-Gln | ChEBI ID-144461 | PubChem CID-71471965 | -6.2 |
|  | Glu-Glu-Lys | ChEBI ID-144461 | PubChem CID-11441068 | -5.3 |
|  | Gly-Cys-Leu | ChEBI ID-144458 | PubChem CID-145455600 | -4.8 |
| HyHEL-63<br>Complexed with<br>HEL mutant<br>K96A<br>(PDB ID: 1NBY) | Arg-His-Trp | ChEBI ID-152557 | PubChem CID-145453842 | -6.9 |
|  | Pro-Ile-Glu | ChEBI ID-144613 | PubChem CID-145457492 | -6.1 |
|  | Cys-Gln-Gln | ChEBI ID-144461 | PubChem CID-71471965 | -5.7 |
|  | Glu-Glu-Lys | ChEBI ID-144461 | PubChem CID-11441068 | -5.9 |
|  | Gly-Cys-Leu | ChEBI ID-144458 | PubChem CID-145455600 | -5.3 |
| Extracellular<br>domain of Pep T1<br>(PDB ID: 5A9D) | Arg-His-Trp | ChEBI ID-152557 | PubChem CID-145453842 | -6.4 |
|  | Pro-Ile-Glu | ChEBI ID-144613 | PubChem CID-145457492 | -5.9 |
|  | Cys-Gln-Gln | ChEBI ID-144461 | PubChem CID-71471965 | -5.8 |
|  | Glu-Glu-Lys | ChEBI ID-144461 | PubChem CID-11441068 | -5.5 |
|  | Gly-Cys-Leu | ChEBI ID-144458 | PubChem CID-145455600 | -5.2 |
| Phosphorylated<br>Map Kinase<br>ERK2<br>(PDB ID: 2ERK) | Arg-His-Trp | ChEBI ID-152557 | PubChem CID-145453842 | -7.9 |
|  | Pro-Ile-Glu | ChEBI ID-144613 | PubChem CID-145457492 | -6.1 |
|  | Cys-Gln-Gln | ChEBI ID-144461 | PubChem CID-71471965 | -5.7 |
|  | Glu-Glu-Lys | ChEBI ID-144461 | PubChem CID-11441068 | -6.0 |
|  | Gly-Cys-Leu | ChEBI ID-144458 | PubChem CID-145455600 | -5.7 |
| 2RH1<br>Beta2-adrenergic<br>G protein-coupled<br>receptor<br>(PDB ID: 2RH1) | Arg-His-Trp | ChEBI ID-152557 | PubChem CID-145453842 | **-8.6** |
|  | Pro-Ile-Glu | ChEBI ID-144613 | PubChem CID-145457492 | **-7.0** |
|  | Cys-Gln-Gln | ChEBI ID-144461 | PubChem CID-71471965 | -5.7 |
|  | Glu-Glu-Lys | ChEBI ID-144461 | PubChem CID-11441068 | -6.3 |
|  | Gly-Cys-Leu | ChEBI ID-144458 | PubChem CID-145455600 | -5.7 |
|  | Isoproterenol<br>(A Known<br>Agonist) | ChEBI ID-64317 | PubChem CID-5808 | **-7.0** |

### Molecular docking analysis

Identifications of these intracellular tripeptides derived of FBS raised various questions on the biological role of these tripeptides in MCF-7 cancer cells. We used initial screening on molecular binding affinity of these tripeptides with various extracellular and intracellular progrowth receptors and proteins. A representative screening of the molecular blind docking data suggested us to look into the potential molecular interactions with β2AR that showed highest binding affinity (−8.6 Kcal/mol) (Table 1). Furthermore, site specific docking on β2AR with Arg-His-Trp (−9.8 Kcal/mol) suggested a favorable binding affinity that is competitive and even better than a known agonist Isoproterenol (−7.0 Kcal/mol) (Table 2). Among all identified intracellular tripeptides, binding affinity is reported as Arg-His-Trp (−9.8 Kcal/mol) > Pro-Ile-Glu (−7.7 Kcal/mol) > Pro-Ile-Glu (−7.7 Kcal/mol) which is less than that of a known agonist Isoproterenol (−7.0 Kcal/mol) > Cys-Gln-Gln (−6.8 Kcal/mol) > Glu-Glu-Lys (−6.3 Kcal/mol) > Gly-Cys-Leu (−6.1 Kcal/mol). Besides the docking affinity, intermolecular interaction analysis suggested that the novel Arg-His-Trp displayed key residues including Thr-195, Phe-194, Ala-200, Lys-305, Tyr-308, Ile-309, Asp-300, Phe-193, His-296, Asn-293 with 09 non-covalent interactions including conventional hydrogen bonds (Figure 4 A, B, C and D). While Asn-312, Asp-113, Thr-118, Phe-193, Phe-290, Val-114 and Phe-289 residues of β2AR are indicted to establish 06 bonds against a known β2AR agonist Isoproterenol (Figure 5A, B, C and D).

**Figure 4.**
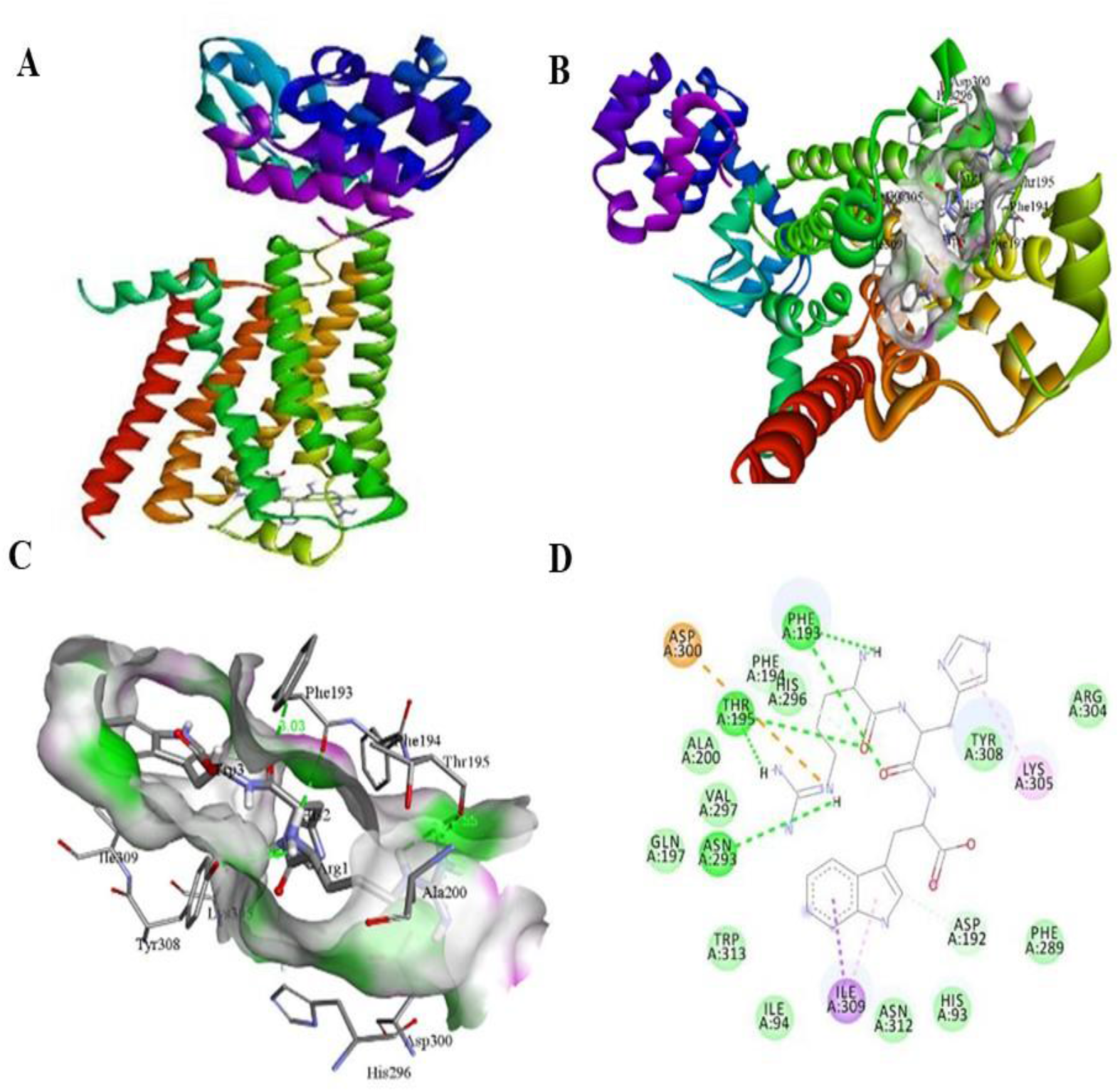
The intracellular tripeptide Arg-His-Trp in MCF-7 cancer cells shows strong molecular binding with β2AR. Molecular interactions between Arg-His-Trp and β2AR were determined in silico with the help of AutoDock Vina and Discovery Studio Visualizer. (A). A ribbon structure with full 3D view between Arg-His-Trp and β2AR. (B). An emphasized 3-D ribbon structure on docked complex between Arg-His-Trp and β2AR. (C). Discovery Studio Visualizer assisted 3-D image of docked molecular structure between Arg-His-Trp and β2AR showing H-bond interaction (Green colour) and steric interaction (Violet colour) between ligand and target protein. (D). Discovery Studio Visualizer assisted 2-D image of docked molecular structure between Arg-His-Trp and β2AR.

**Figure 5.**
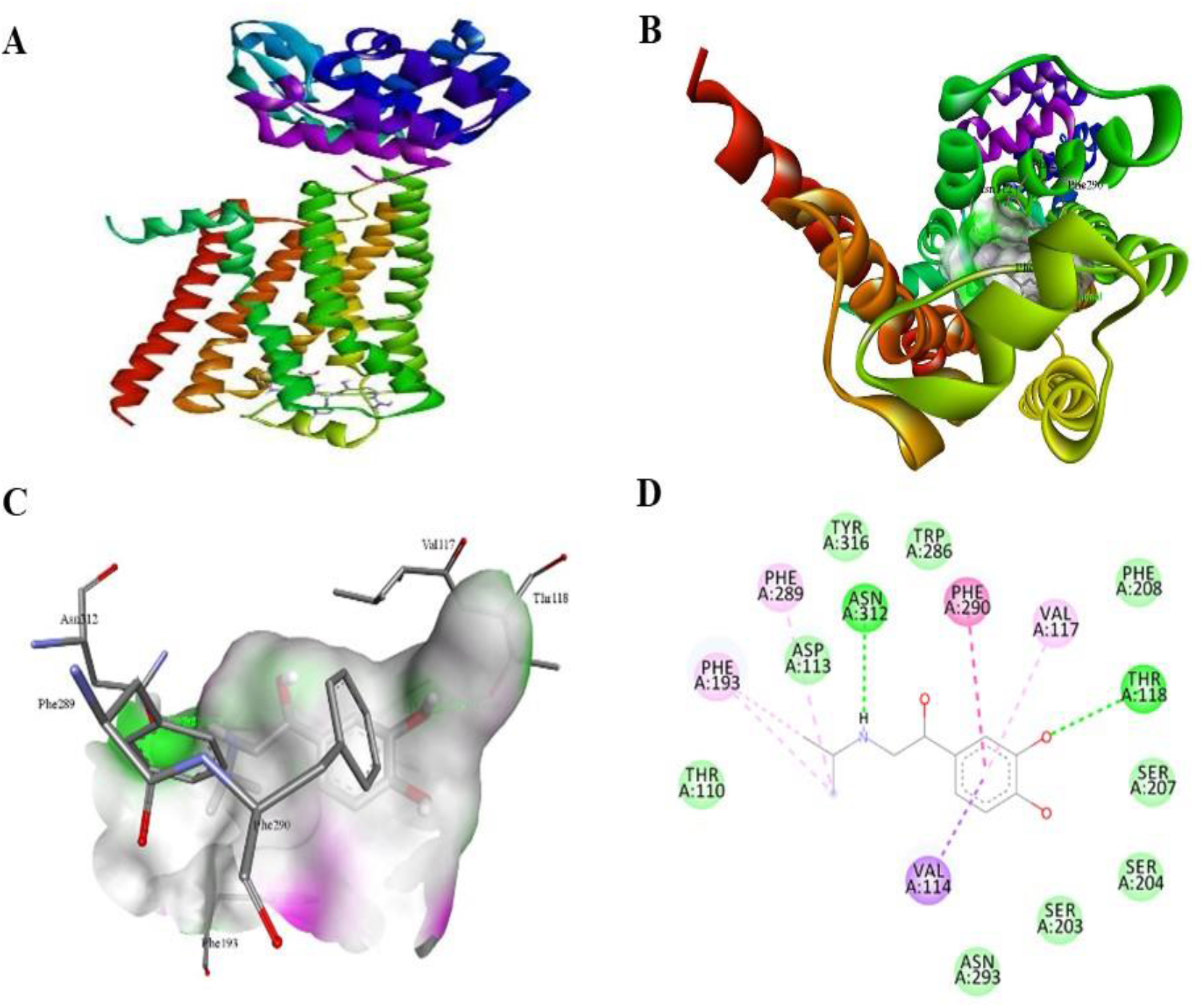
A known agonist isoproterenol shows agonistic binding affinity β2AR. Molecular interactions between isoproterenol and β2AR were determined with the help of AutoDock Vina and Discovery Studio Visualizer. (A). A secondary structure of receptor in ribbon structure with full 3D view between isoproterenol and β2AR. (B). An emphasized 3-D ribbon structure on docked complex between isoproterenol and β2AR. (C). Discovery Studio Visualizer assisted 3-D image of docked molecular structure between isoproterenol and β2AR showing H-bond interaction (Green colour) and steric interaction (Violet colour) between ligand and target protein. (D). Discovery Studio Visualizer assisted 2-D image of docked molecular structure between isoproterenol and β2AR.

**Table 2:** The details on site specific binding affinities of identified tripeptides from FBS with Beta-2 Adrenergic receptor (PDB ID: 2RH1). Here, molecular interactions between tripeptides and a known agonist are evaluated by using AutoDock Vina and visualized by the help of Discovery Studio Visualizer that reveals the nature of amino acid residues, number of bonds, nature of bonds and distance of bonds.

| Name of Ligand and its PubChem CID | Binding affinity (kcal/mol) | Binding Residues | No. of Bonds | Bond Distances (Å) |
| --- | --- | --- | --- | --- |
| Arg-His-Trp<br>(PubChem CID-145453842) | -9.8 | Thr-195, Phe-194, Ala-200, Lys-305, Tyr-308, Ile-309, Asp-300, Phe-193, His-296, Asn-293. | 09 | 2.91, 4.90, 4.61, 3.86, 3.46, 3.03, 3.75, 4.00, 4.21. |
| Pro-Ile-Glu<br>(PubChem CID-145457492) | -7.7 | Tyr-308, Asn-293, Thr-195, Tyr-199, Phe-194, Phe-193 | 07 | 4.09, 4.65, 4.98, 5.05, 3.15, 3.63, 3.05 |
| Cys-Gln-Gln<br>(PubChem CID-71471965) | -6.8 | Lys-305, Asn-301, Asp-300, Asn-293, Ala-200, Tyr-199, Phe-193, Thr-195, Gln-2, Gln-3. | 09 | 2.29, 2.32, 5.33, 5.66, 5.16, 2.74, 2.71, 2.02, 3.15. |
| Glu-Glu-Lys<br>(PubChem CID-11441068) | -6.3 | Phe-322, Ser-329, Arg-131, Thr-66, Asn-69, Leu-275, Ala-271, Lys-1, Glu-2, Glu-3. | 07 | 2.97, 2.98, 3.00, 3.59, 5.31, 3.03, 2.78. |
| Gly-Cys-Leu<br>(PubChem CID-145455600) | -6.1 | Ala-271, Tyr-141, Ser-143, Arg-131, Asn-69, Thr-68, Val-67, Gln-1, Cys-2, Cys-3. | 10 | 2.20, 4.60, 2.90, 2.64, 2.67, 2.26, 2.51, 3.16, 2.99, 3.27. |
| Isoproterenol<br>(PubChem CID 5808) (Known Agonist) | -7.0 | Asn-312, Asp-113, Thr-118, Phe-193, Phe-290, Val-114, Phe-289 | 06 | 2.2, 3.2, 5.45, 4.89, 4.97, 4.5 |

### Molecular dynamics simulation

Furthermore, we performed molecular dynamics simulations of Arg-His-Trp-(β2AR) complex to confirm its binding affinity with β2AR and binding duration. A Root Mean Square Deviation (RMSD) graph reveals that alignment of the ligand-protein complex with protein backbone (C-α atoms) occurs strongly and constantly up to 10 ns (Figure 6). In fact, after 6 ns of simulation, stability of Arg-His-Trp-(β2AR) becomes even better. By analyzing the RMSD plot of Arg-His-Trp and β2AR, a difference in the range of less than 1 Å was observed which is better than the acceptable range of 1-3 Å that indicates an efficient stability. Interestingly, Arg-His-Trp-(β2AR) contact graph showed the contribution of interacting key residues namely Thr-195, Asp-113, Phe-193, Asn-293, His-296, Tyr-308 and Asn-312 to create stable interacting bonds which are mainly hydrogen bonds (Figure 7A and B).

**Figure 6.**
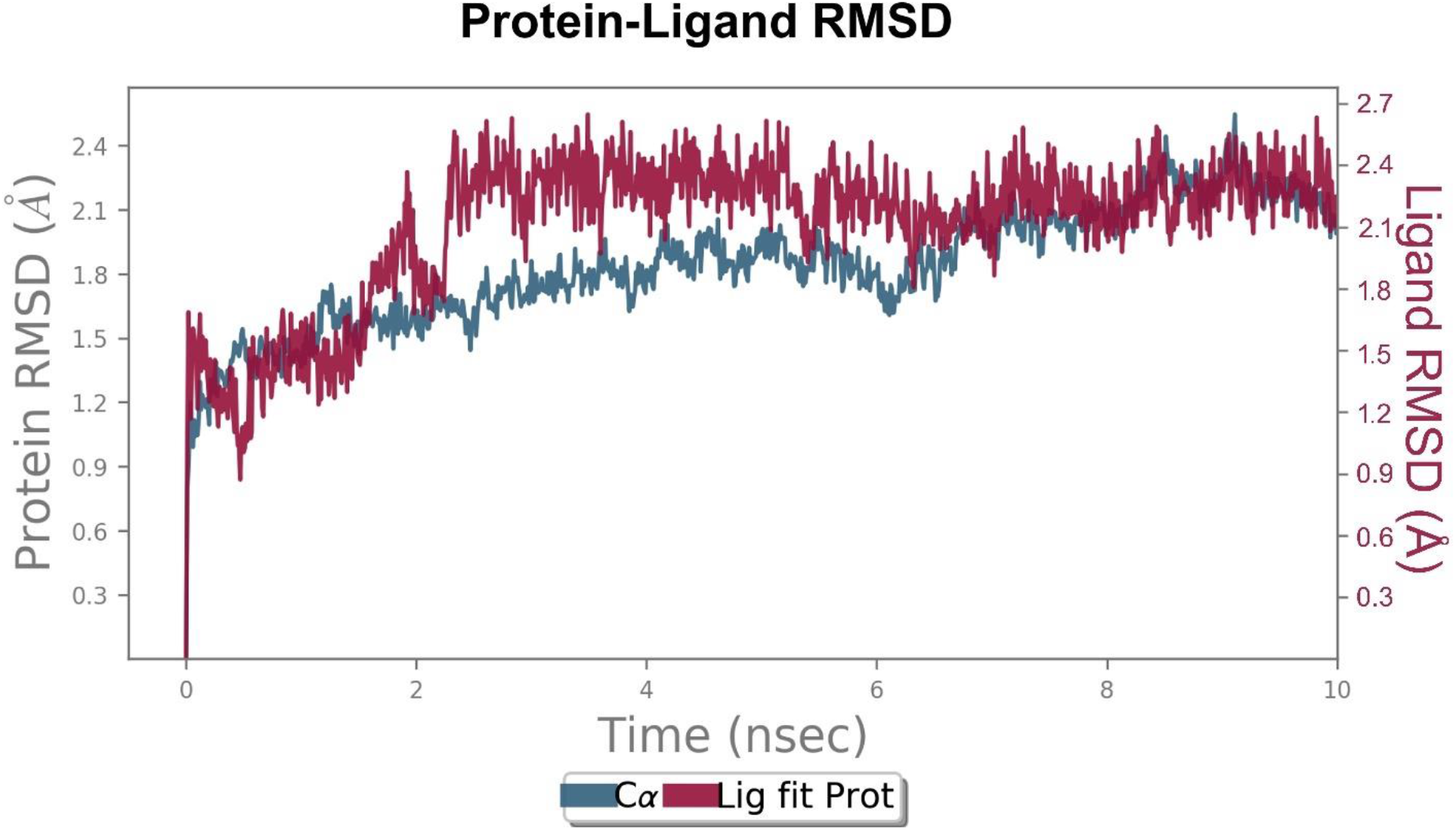
Binding between Arg-His-Trp and β2AR is in stable alignment by molecular dynamics and simulations. Protein-ligand RMSD plot of Arg-His-Trp and β2AR generated by Molecular Dynamics Simulations.

**Figure 7.**
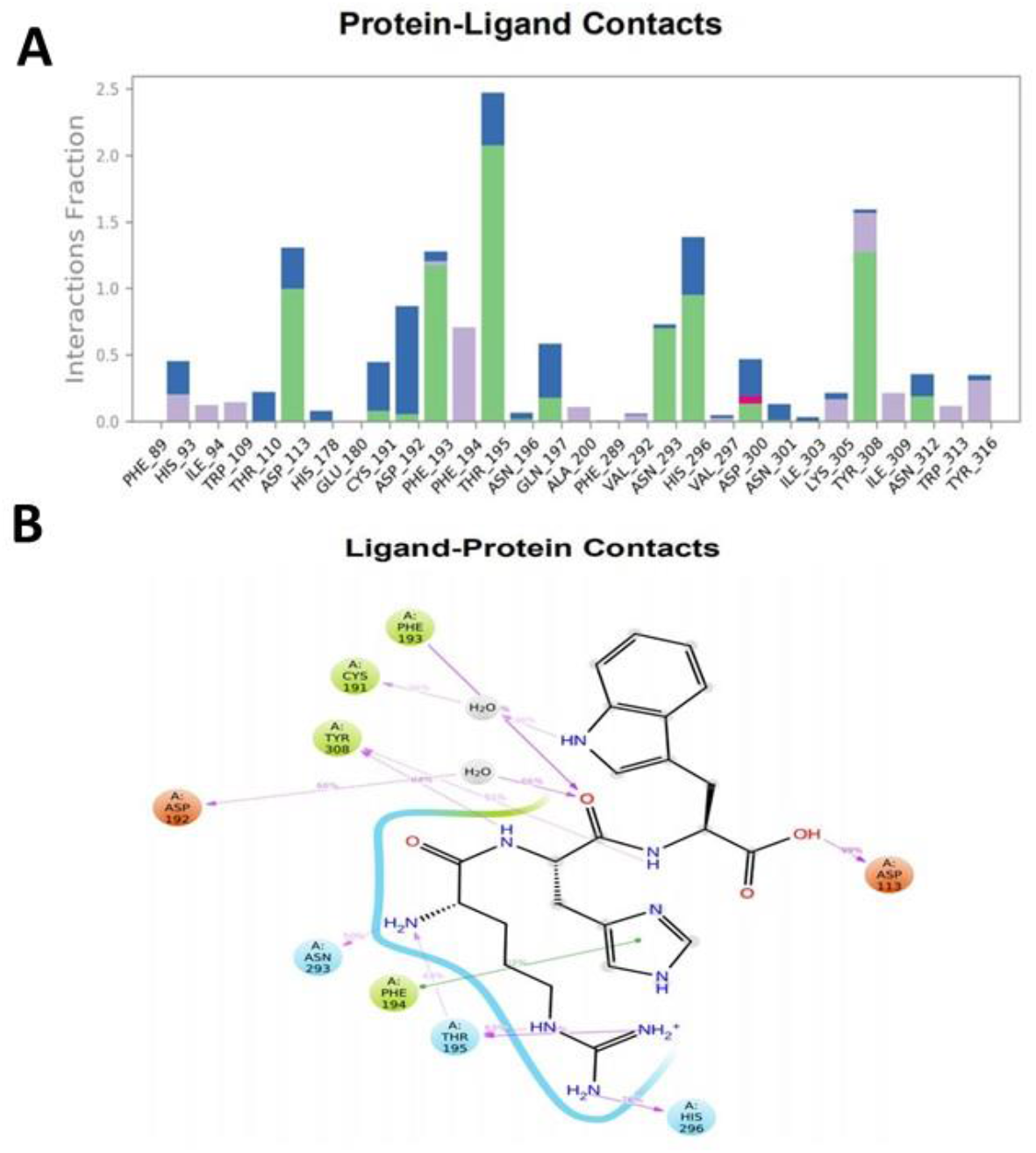
Arg-His-Trp displays a strong and specific contact with amino acid residues that contribute to agonistic effects in β2AR. **(A)** Arg-His-Trp and β2AR contact generated by molecular dynamics simulations. **(B)** A detailed ligand atom interaction with the agonistic effects contributing amino acid residues of β2AR.

## Discussion

In view of available data and need to explore new classes of agonists to β2AR, appreciable attempts are in progress. It is well-known that during in vitro culture of breast cancer cells such as MCF-7 cells, along with required media, FBS is required to augment the growth and proliferation needs of the growing cells. In literature, FBS is known to contain various organic and inorganic components including ions besides albumin, growth factors and many more pro-growth and proliferation factors. However, there is a gap in the attempts to know the nature of small compounds such as tripeptides from FBS that have a potential role in growth and proliferation of in vitro grown cells. Therefore, we asked a basic question on the possibilities of a tripeptide mimetic as an agonist of β2AR from FBS that supports the growth of cancer cells. Interestingly, there has been no attempt to explore the nature of agonists from FBS in the form of tripeptide mimetic so far.

We believe that non-availability of data on the agonists from FBS may be attributed to the complex nature of FBS and it is difficult to study the intracellular components which have entered from external media such as FBS. In this direction, a reported novel vertical tube gel electrophoresis (VTGE) approach from our lab appears to be suitable which will help characterizing small size molecules including tripeptides of FBS and at the same time excluding large size components such as growth factors, albumin etc.

As expected in line with earlier reports, MCF-7 cells showed desired optimal growth and viability in the presence of FBS. Hence, interpretation is drawn that components of FBS including growth factors, cytokines and additional potential factors are responsible for such effects. However, additional factors in the form of tripeptides of FBS origin is not reported. Therefore, identifications of these novel intracellular tripeptides namely Arg-His-Trp, Pro-Ile-Glu, Cys-Gln-Gln, Glu-Glu-Lys, and Gly-Cys-Leu prompted us to explore their potential contribution in the growth and proliferation of MCF-7 cancer cells. Furthermore, data indicated that novel intracellular tripeptides including most prominent Arg-His-Trp is highly effective in binding to β2AR and this receptor is involved in the growth and proliferation of breast cancer cells. However, literature draws some ambiguity in terms of role of agonists and antagonists of β2AR in different molecular sub-types of breast cancer cells.

Therefore, the role of agonists of β2AR in growth and proliferation of breast cancer cells is determined by the molecular subtypes and expressions of molecular markers. Furthermore, agonists such as norepinephrine and isoproterenol have been reported to promote the growth, proliferation, invasiveness, and metastasis of breast cancer cells specifically hormone and HER positive subtypes (Pérez Piñero et al., 2012). Conversely, antagonists are shown to arrest the growth and proliferation of these breast cancer cells. Besides the known agonists and antagonists of β2AR, there is a promising avenue to explore new class of pharmacological inhibitors and peptide mimetic that can work as agonists or antagonists and explore for their role in human diseases. In this paper, we suggest that Arg-His-Trp with potential agonistic effects upon β2AR may contribute to the growth and proliferation of MCF-7 breast cancer cells and such possibilities are expected.

In our study, the most efficient intracellular tripeptide Arg-His-Trp displays a binding affinity even better as compared to isoproterenol, a known agonist of β2AR. Besides binding affinity, detailed analysis of molecular interactions predicted the stable and efficient nature of bonds compared to isoproterenol. Interestingly, molecular docking and molecular dynamics simulations data clearly shows that key residues including Thr-195, Asp-113, Phe-193, Asn-293, His-296, Tyr-308 and Asn-312 are responsible for high binding affinity and stabilization of Arg-His-Trp with β2AR and such binding patterns are similar to earlier reports on key residues that contribute to agonistic effects.

Similar to our molecular docking data, molecular dynamics simulations and crystallography studies revealed the binding sites (Asp-113, Asn-312, Phe-193) of an agonist isoprenaline upon β2AR involving various non-covalent interactive forces including hydrogen, disulfide and non-polar bonds (Vanni et al., 2011; Rosenbaum et al., 2011). An interesting evidence suggests that several ligands may display as potential agonists, because β2AR displays dynamic conformation changes. Most of agonists are endowed with distinct interacting patterns that stabilize β2AR during their agonistic effects (Kahsai et al., 2011). Key amino acid residue Asn-293 located in the upper half of transmembrane helix VI is shown to be substantial for the agonistic effects of drug against β2AR (Wieland et al., 1996).

In literature, β2AR is known as one of the highly studied GPCR family of receptors in view of structure and function determined by in silico, molecular dynamics, crystallography, and various other molecular biology tools. Based on all such studies, several types of natural and artificial agonists are reported, most notably epinephrine and isoproterenol. These agonists are suggested to bind to specific amino acid residues (Asp-193, Thr-195, Phe-193, Asn-312, Asp113, Tyr-308) that activate the function of GPCRs and finally the level of cAMP is elevated that helps in the modulation of various cellular function including proliferation. Based on the existing views and data collected from the present study, a potential agonistic role of this novel tripeptide Arg-His-Trp is proposed against β2AR.

An obvious concern on toxicity is raised for any novel agonists or antagonists as drugs of any human disease conditions. Here, we defend that the novel tripeptide Arg-His-Trp is of biological source (FBS origin) and hence the chance of toxicity or side effects will be minimal. In a potential absorption, distribution, metabolism, excretion, and toxicity (ADMET) prediction model (Schyman et al., 2017), our data is appreciable to show the complete lack of cytotoxicity, mutagenicity, cardiotoxicity, drug-drug interactions, microsomal stability, and drug-induced liver injury by Arg-His-Trp as compared to other known drugs including isoproterenol (Figure S6). Here, maximum recommended therapeutic dose (MRTD) of Arg-His-Trp (14617 mg/ daily dose) is highly desirable over isoproterenol, the known agonist of β2AR.

There is evidence on tripeptides from natural sources including FBS and plasma or synthetic sources for their various biological roles including pro-growth, antioxidant defense and differentiation (Sensenbrenner et al., 1975). It is conceivable that FBS is a good sources of growth factors and other biological components including tripeptides. At the same time, all tripeptides may not serve the same molecular effects for the growth and proliferation of growing cells. This is well supported from our observations that all detected intracellular tripeptides did not show efficient binding affinity with β2AR. In this context, among identified intracellular tripeptides, Glu-Glu-Lys (binding affinity −6.3 Kcal/mol) and residues Phe-322, Ser-329, Arg-131, Thr-66, Asn-69, Leu-275, Ala-271, Lys-1, Glu-2, Glu-3) and Gly-Cys-Leu (−6.1 Kcal/mol and residues Ala-271, Tyr-141, Ser-143, Arg-131, Asn-69, Thr-68, Val-67, Gln-1, Cys-2, Cys-3) are not agonists of β2AR (2RH1). Hence, these two tripeptides may be assigned different biological functions. On the other hand, other three tripeptides and most notably Arg-His-Trp is predicated to be a strong agonist of β2AR.

Taken together, this paper reports on tripeptides as potential agonists of β2AR by using VTGE approach and molecular docking and dynamics on a model MCF-7 cells those are grown in the presence of routinely used culture media and FBS components. In future, agonistic effects of Arg-His-Trp may be translated into its therapeutic use against various ailments including cardiovascular diseases. A model on agonistic effect of intracellular tripeptide is depicted (Figure 8).

**Figure 8.**
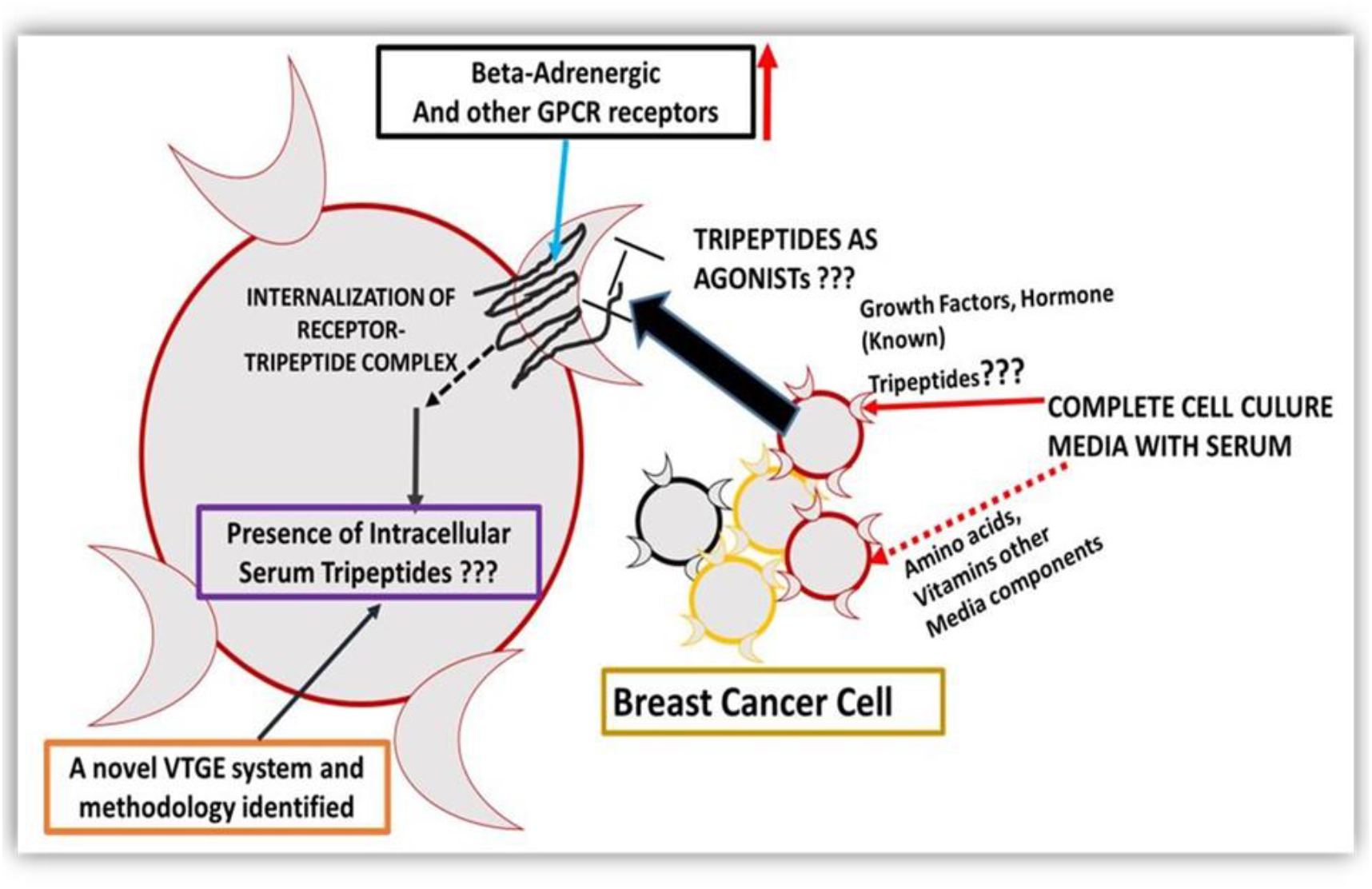
A proposed model on the agonistic effect of intracellular tripeptide against β2AR.

## Conclusion

In the literature, limited data exists that suggests the presence of tripeptides in FBS and other biological sources including urine for various biological properties. In our studies, we report on the biological relevance of a novel intracellular tripeptide that was derived from FBS in case of growing MCF-7 cancer cells. Furthermore, with the help of molecular docking and molecular dynamics and simulations tools, we show that the most prominent tripeptide Arg-His-Trp has an agonistic effect upon β2AR. This novel tripeptide appears to show a better agonistic effect, and minimal side effects as compared to other known agonists of β2AR. Therefore, the present findings suggest a futuristic approach to explore the tripeptide Arg-His-Trp as an agonist of β2AR and its therapeutic applications in cardiovascular diseases.

---

**Figure S1.**
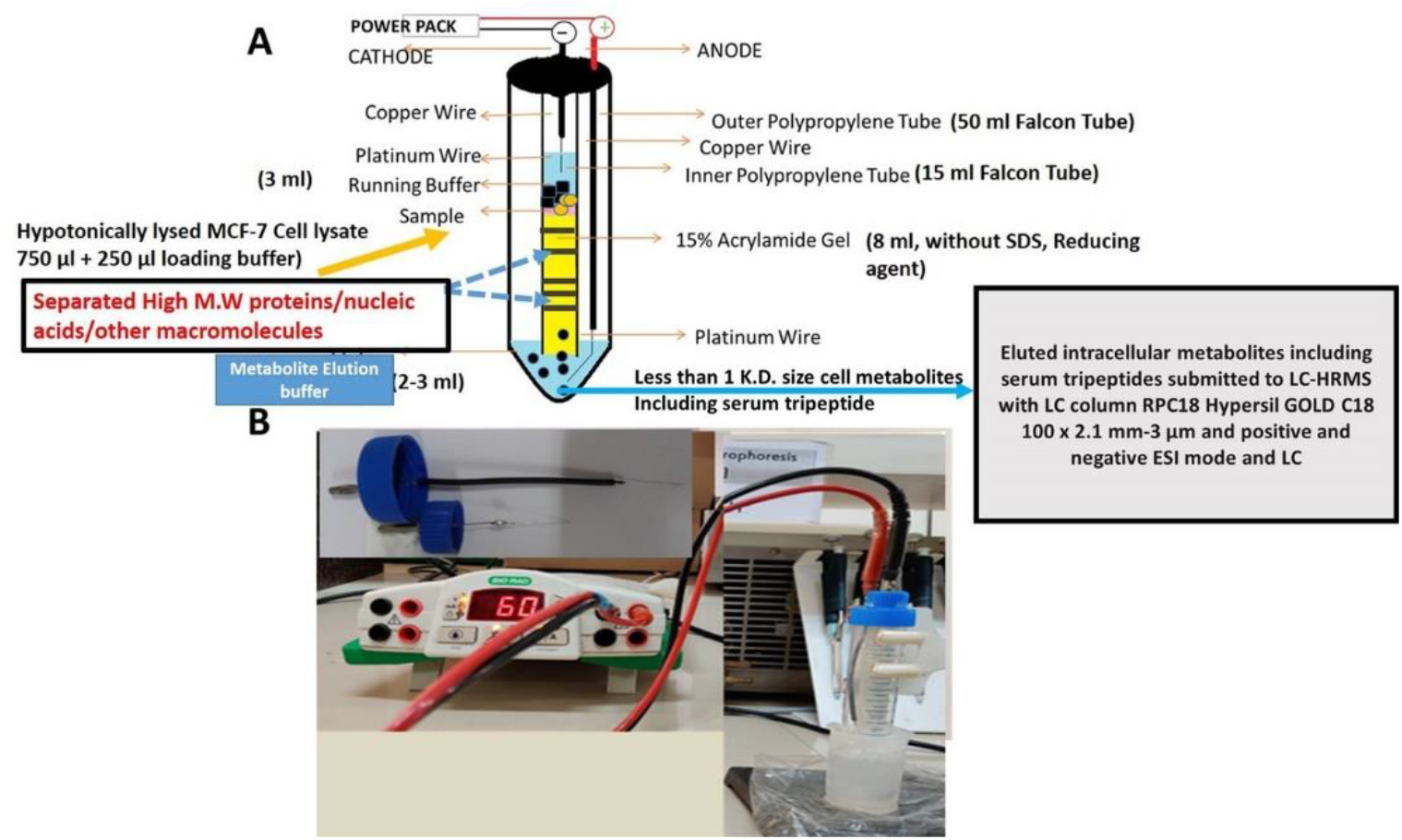
Model of a novel vertical tube gel electrophoresis (VTGE) metabolite purification system. (A) A flow diagram of VTGE system with details on matrix and buffer. (B) A photograph of VTGE system in a running condition.

**Figure S2.**
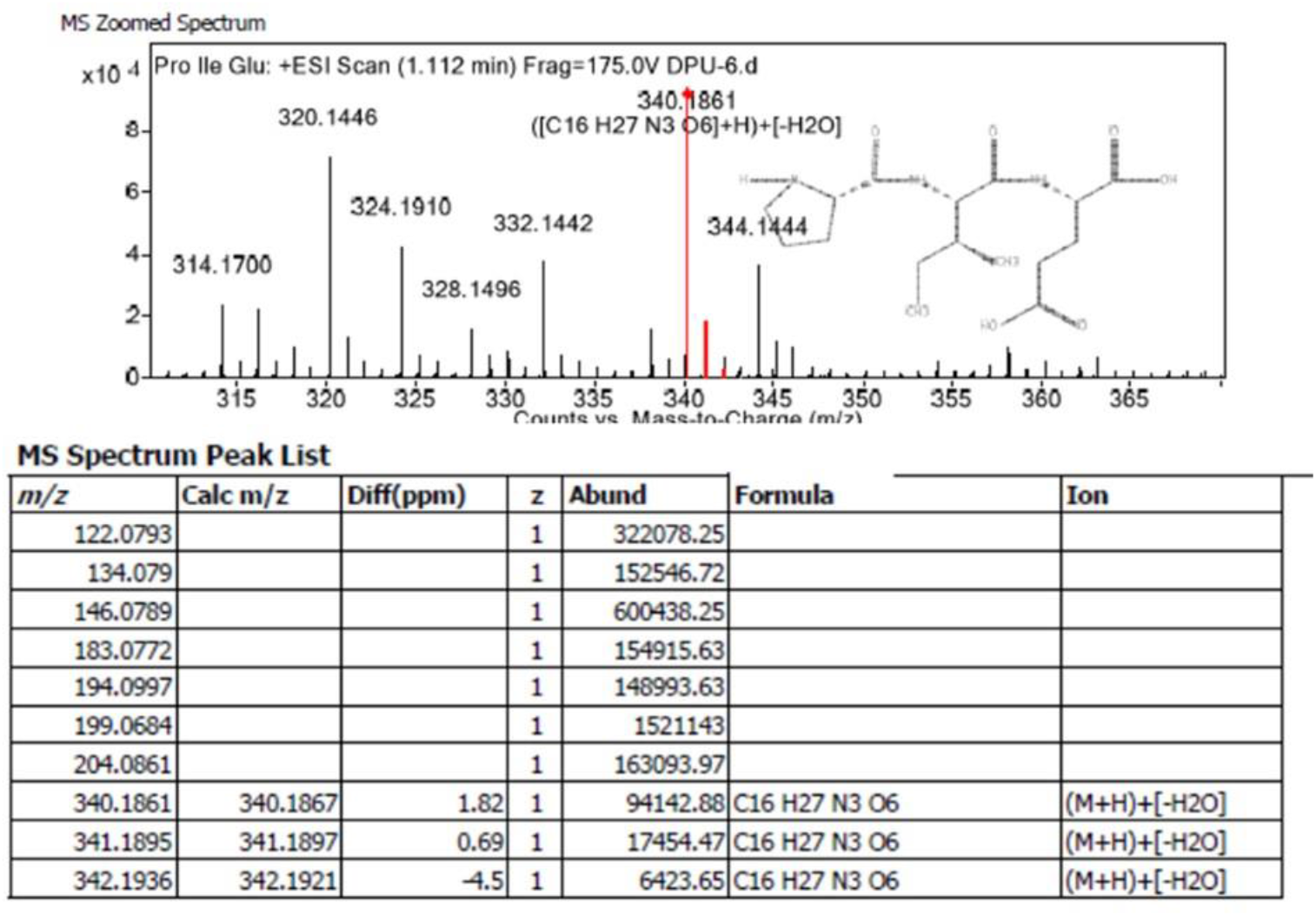
An intracellular tripeptide Pro-Ile-Glu of FBS origin is detected MCF-7 cancer cells. A positive ESI-MS spectrum along with peak lists of Pro-Ile-Glu.

**Figure S3.**
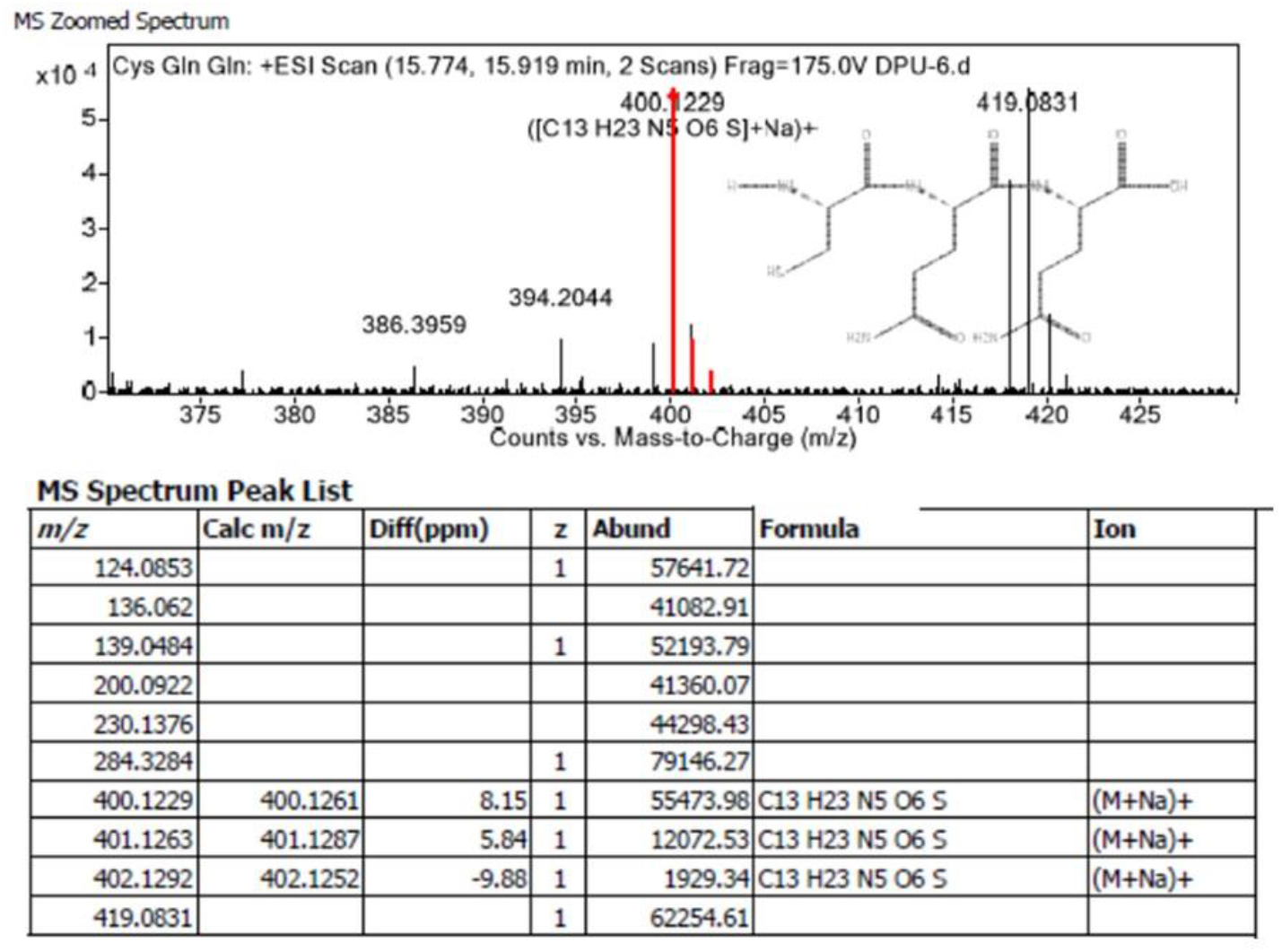
An intracellular tripeptide Cys-Gln-Gln of FBS origin is detected MCF-7 cancer cells. A positive ESI-MS spectrum along with peak lists of Cys-Gln-Gln were detected by LC-HRMS.

**Figure S4.**
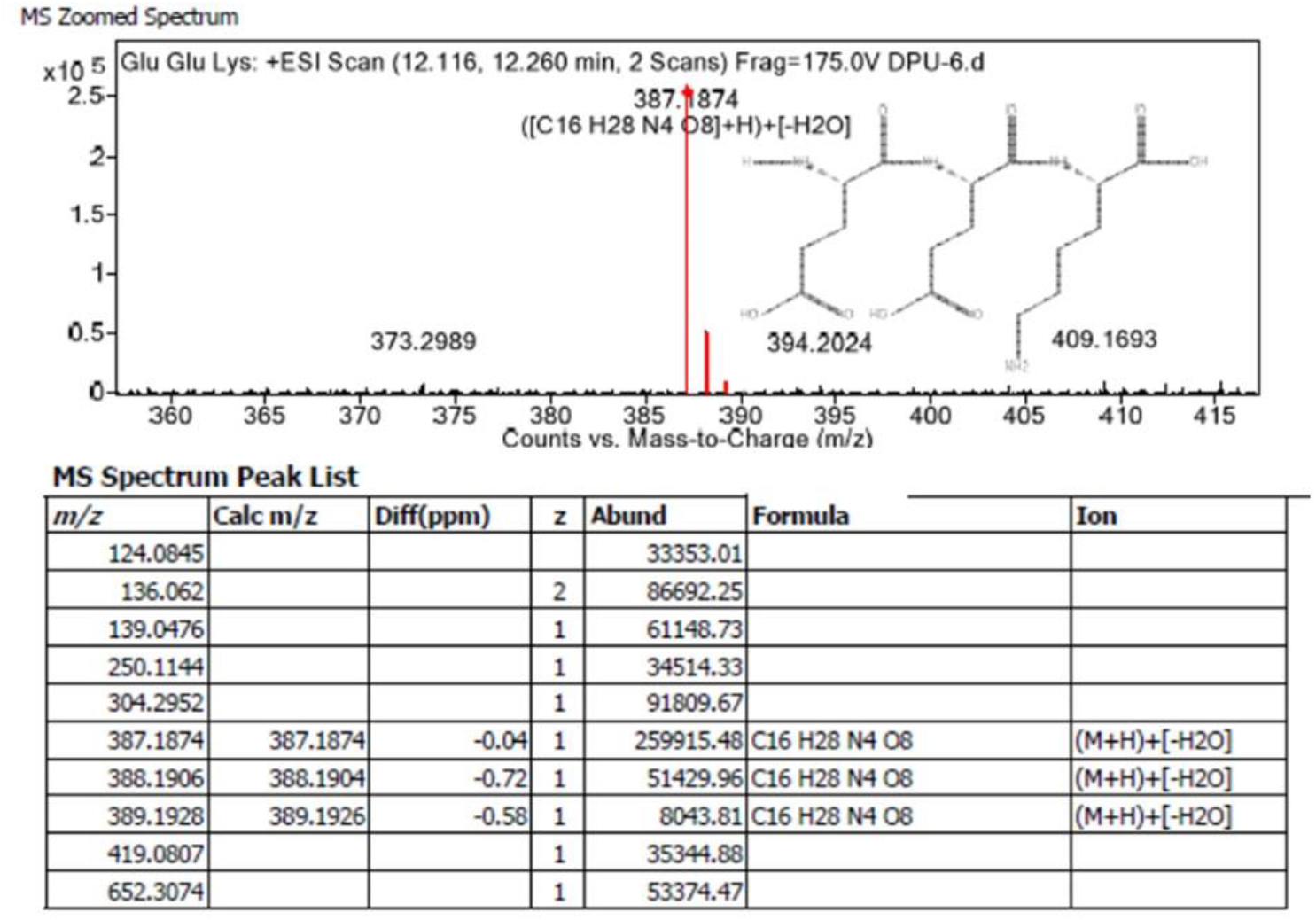
An intracellular tripeptide Glu-Glu-Lys of FBS origin is detected MCF-7 cancer cells. A positive ESI MS spectrum with specific peak lists of Glu-Glu-Lys were detected by LC-HRMS.

**Figure S5.**
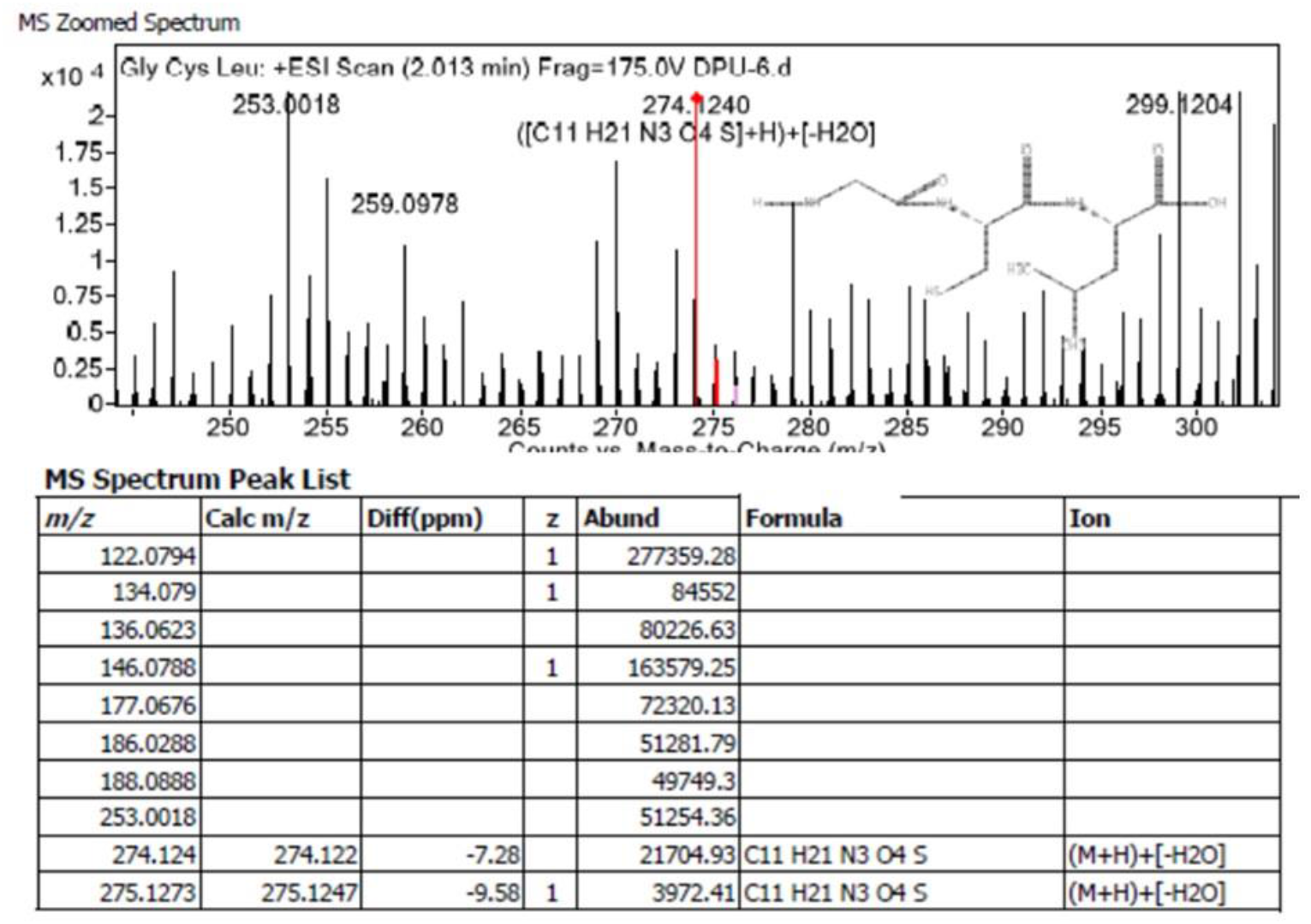
An intracellular tripeptide Gly-Cys-Leu of FBS is detected MCF-7 cancer cells. A positive ESI-MS spectrum along with specific peak lists of Gly-Cys-Leu were detected by LC-HRMS.

**Figure S6.**
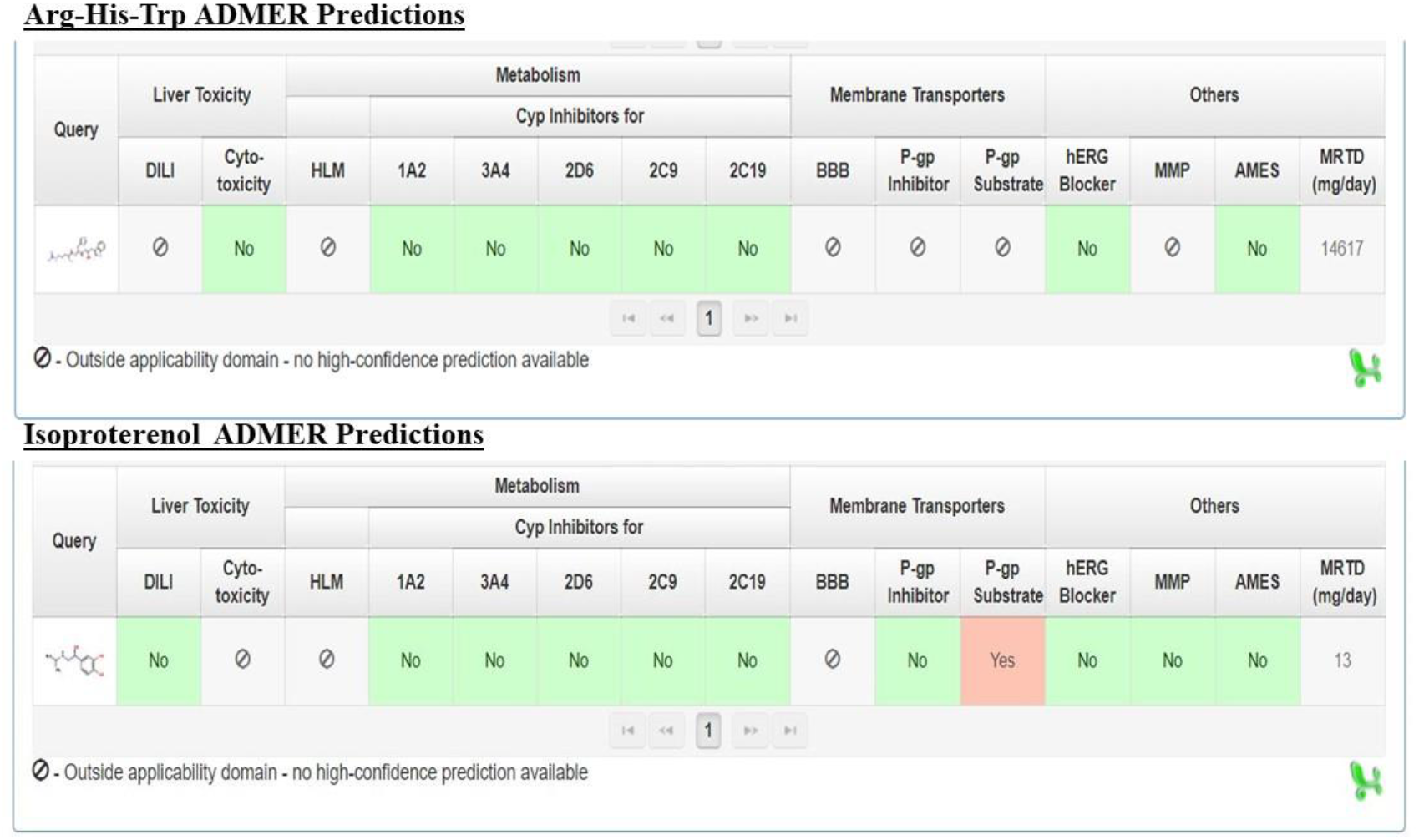
A novel tripeptide Arg-His-Trp is predicted for minimal toxicity and tissue injury by ADMET prediction tool.

